# Adrenal extramedullary hematopoiesis as an inducible model of the adult hematopoietic niche

**DOI:** 10.1101/2023.03.15.531679

**Authors:** Frédérica Schyrr, Alejandro Alonso-Calleja, Anjali Vijaykumar, Sandra Gebhard, Rita Sarkis, Silvia F. Lopes, Aurélien Oggier, Laurence De Leval, César Nombela-Arrieta, Olaia Naveiras

## Abstract

Hematopoietic Stem and Progenitor Cells (HSPCs) reside in the hematopoietic niche, a structure that regulates the balance of cellular quiescence, self-renewal and commitment in a demand-adapted manner. The bone marrow (BM) hematopoietic niche is formed by several cellular players, mainly endothelial cells, osteoblasts, adipocytes, and stromal cells. While the BM niche forms a complex structure, evidence exists for simpler, albeit functional, extramedullary hematopoietic niches. However, the composition of what constitutes the simplest unit of an HSPC supportive microenvironment remains largely unknown. Here, we show that the adult adrenal gland can be transformed into a hematopoietic supportive environment. Upon splenectomy and hormonal stimulation, the adult adrenal gland can be induced to recruit and host HSPC function, including serial transplantation. Furthermore, the adrenal stroma contains a CXCL12+ population, reminiscent of BM CXCL12-Abundant Reticular (CAR) cells. Mirroring this, we found CXCL12+ cells in patient samples obtained from a local cohort of myelolipoma, a benign adrenal tumor composed of adipose and hematopoietic tissue that constitutes the most common site of extramedullary hematopoiesis specific to the adult. We present our model as a novel tool to increase our understanding of the physiology of hematopoietic support and to facilitate the development of a boneless niche model.

## Introduction

Hematopoietic Stem and Progenitor Cells (HSPCs) reside in niches within the bone marrow (BM), structures that regulate the balance of cellular quiescence, self-renewal and commitment towards progenitor cells in a demand-adapted manner. This tightly regulated process is controlled by a combined input of different soluble factors, such as cytokines, and non-soluble cues that require cell-cell contact. The hematopoietic niche is formed by several cellular players, mainly endothelial cells, osteoblasts, adipocytes, and a variety of stromal progenitor cells (1–3).

Perturbations of the BM stromal cell (BMSC) compartment have been described to impact hematopoiesis (4, 5) and to drive leukemic transformation (6–8). This has spurred efforts to develop reliable models to study the stromal component of the BM niche, a goal that has proven to be challenging both in vitro (9) and in vivo (10–14). The standards of the field include the formation of subcutaneous ossicles that contain hematopoietic tissue through the co-injection of BMSCs and a carrier material (like bone powder or Matrigel®), often together with daily injections of parathyroid hormone to induce ossification of the nodules, which occurs in 8 to 12 weeks (10, 11). Consequently, current methods of producing ossified nodules for BM modelling are both extremely time-consuming and limited by their unpredictable efficiency to form ossified heterotopic BM, partly due to batch-to-batch variability of the biological carriers (15).

While adult hematopoiesis takes place primarily in the BM, examples of adult hematopoiesis outside bones exist, grouped under the term extramedullary hematopoiesis (EMH). In both mice and human, EMH can be classified in two groups (16) (i) EMH arising in fetal hematopoiesis sites, fundamentally spleen and liver, and (ii) EMH in non-fetal hematopoietic sites, typically paraspinal and peritoneal locations, where the hematopoietic tissue is embedded within adipocytic tissue, which we refer to as adipose-associated EMH. While white adipose tissue has been shown to host functional HSCs in a quiescent state (17, 18), active proliferation of hematopoietic progenitors within the adipose tissue occurs only in the context of adipose-associated EMH.

Adipose-associated EMH can arise either spontaneously, which is considered a benign tumor possibly driven by a stromal population (19, 20), or upon extreme hematopoietic demand, typically in patients suffering from thalassemia (21). Benign adipose-associated EMH occurs with a particular high frequency in the adrenal gland, constituting a distinct clinical entity called adrenal myelolipoma, with a prevalence at autopsy estimated to be around 0.08 to 0.2% (20). Of note, the vast majority of adrenal myelolipomas are unilateral and sporadic. Bilateral cases are associated with congenital adrenal hyperplasia, an autosomal recessive disorder characterized by a defect (partial or total) in the enzymatic pathways of the adrenal cortex, most frequently affecting the enzyme steroid 21-hydroxylase. The defective function of the adrenal gland in the context of congenital adrenal hyperplasia leads to a decrease in the production of cortisol and aldosterone. The decrease in cortisol levels results in an increase of adrenocorticotropic hormone (ACTH), as the production of this hormone is under negative feedback control by cortisol, ultimately leading to adrenal gland hyperplasia (22). Interestingly, the prevalence of myelolipoma in congenital adrenal hyperplasia is estimated to be as high as 6% (23).

We thus hypothesized that exogenous hormonal stimulation may be leveraged to induce a simpler, boneless hematopoietic niche in the adrenal glands. We show that the murine adrenal gland can be induced by ACTH to form a hematopoietic-supportive tissue and used as a model to study the components of a minimalistic hematopoietic niche, thus bypassing the need for an ossified structure. Induced adrenal glands contain serially transplantable HSPCs, host HSPCs delivered after adrenal induction, and retain a CD45+ hematopoietic cells in a CXCR4-dependent manner. Indeed, CXCL12-GFP reporter mice reveal numerous CXCL12+ stromal cells with reticular morphology resembling the so-called CXCL12-Abundant Reticular (CAR). Furthermore, we show that human myelolipoma samples also contain an abundant CXCL12+ population, mirroring the findings of our murine model.

## Results

### The adrenal gland can be hormonally induced to host hematopoietic cells

The challenging complexity of the BM hematopoietic niche prompted us to develop a new model to study the minimal components of the functional hematopoietic niche. In a paper from 1950, Selye and Stone (24) described the possibility of transforming the adrenal gland into myeloid-like tissue with the appropriate hormonal stimulation. Using a combination of pituitary gland extract, sexual and thyroid hormones injected in splenectomized rats, they observed the transformation of the zona fasciculata of the adrenal gland into a structure reminiscent of myeloid tissue. The authors described it as composed of adipocytes and hematopoietic cells, including megakaryocytes, consequently demonstrating *in situ* hematopoiesis in the organ.

Based on this model, we designed a strategy to induce EMH in the murine adrenal gland. We injected a hematopoietic cytokine (Granulocyte colony-stimulating factor (G-CSF)) and the pituitary axis adrenal-corticotropic hormone (ACTH) as well as an androgen (testosterone) in splenectomized C57BL/6J mice (Figure 1A). As expected with the stimulation of the hypothalamic-pituitary-adrenal axis, mice developed, within the first seven days of injection, symptoms of Cushing syndrome, showing increased weight, polyuria and polydipsia (25) (Supplementary Figure 1A-B). EMH induction treatment tended to increase circulatory white blood cells and granulocytes, as expected upon G-CSF administration, but had no effect on hemoglobin (Supplementary Figure 1C-E).

**Figure 1.**
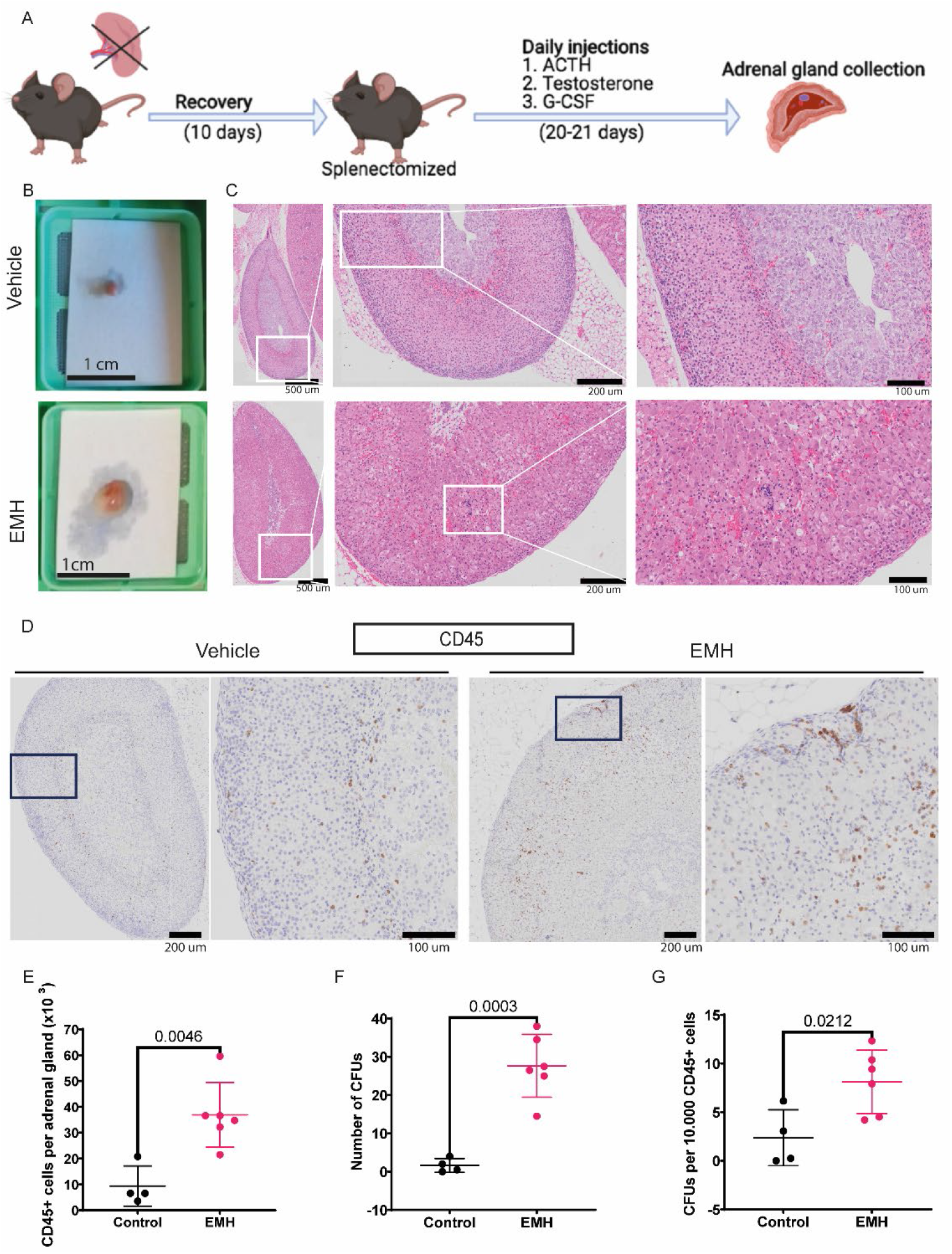
The adrenal gland can be hormonally induced to contain hematopoietic cells. **A** General experimental design to induce EMH in the adrenal gland. **B** Macroscopic picture of freshly isolated adrenal glands. **C** Representative images of H&E stains for vehicle (top) and EMH-induced (bottom) adrenal glands. The boxed area is magnified on the right of each image, dotted circle highlights hematopoietic foci. **D** Representative images of CD45 IHC stain in an adrenal gland. The boxed area is magnified on the right of each image. **E** Number of CD45+ cells per adrenal gland, measured by flow cytometry (control n=4, EMH n=6 mice, two independent experiments). **F** CFU assay, number of colonies per adrenal gland (control n=4, EMH n=6 mice, two independent experiments). **G** CFU assay, number of colonies in the adrenal glands normalized to 10.000 CD45+ cells (control n=4, EMH n=6 mice, two independent experiments). Data are represented as mean ±SD. Differences were assessed using unpaired, two-tailed Student’s *t*-test. P values are indicated in the graphs.

Adrenal glands from induced mice were markedly larger than those from the control group (Figure 1B). Upon histological examination, foci of hematopoietic cells could be identified morphologically in the adrenal cortex in H&E-stained samples (Figure 1C) and further confirmed with IHC for CD45, a pan-hematopoietic marker (Figure 1D). We also found cells positive for vWF, a marker of megakaryocytes, suggesting in situ hematopoiesis (Supplementary Figure 1D) The increased number of hematopoietic cells in the induced adrenal glands was confirmed by flow cytometry (Figure 1E), which showed a 4-fold increase in CD45+ cells in the glands retrieved from the treated group.

Colony-forming unit (CFU) assays measure the progenitor function of short-term HSPCs (26). This assay showed that cells within the induced adrenal glands form more colonies than those obtained from the control glands, with approximately a 15-fold increase (Figure 1F). The increase in CFUs was statistically significant even when normalized to the total CD45+ count to take into consideration the increased adrenal volume and thus higher numbers of CD45+ cells in the induced glands (Figure 1G).

Collectively, these results indicate that the adrenal glands can be hormonally induced to selectively enrich in hematopoietic cells with increased colony-forming potential.

### The induced adrenal gland contains functional, serially transplantable HSPCs

Once we had determined the presence of hematopoietic cells with CFU potential, we investigated the nature of the hematopoietic cells found in the induced adrenal glands. For this we performed flow cytometry for known HSPC surface markers. In the BM, the immunophenotype of the progenitor cells, defined as lineage-, c-Kit+ and Sca-1+ (LKS), showed no difference between control and induced mice (Figure 2A). In blood, the induction cocktail caused an increase in c-Kit+ progenitor cells, as expected due to G-CSF administration. In the adrenal gland, a different surface marker profile was observed. CD45+ lineage-cells obtained from the induced adrenal glands did not show the same surface marker profile as in the BM, but instead a proportion of them displayed a c-Kit^low^ and Sca-1^low^ profile (Figure 2A). Due to the limited number of hematopoietic cells in the adrenal glands and the lower expression of those markers, cell sorting could not be performed to investigate the functional capacities of the lineage-populations present in the induced adrenal glands based on c-Kit or Sca-1 expression.

**Figure 2.**
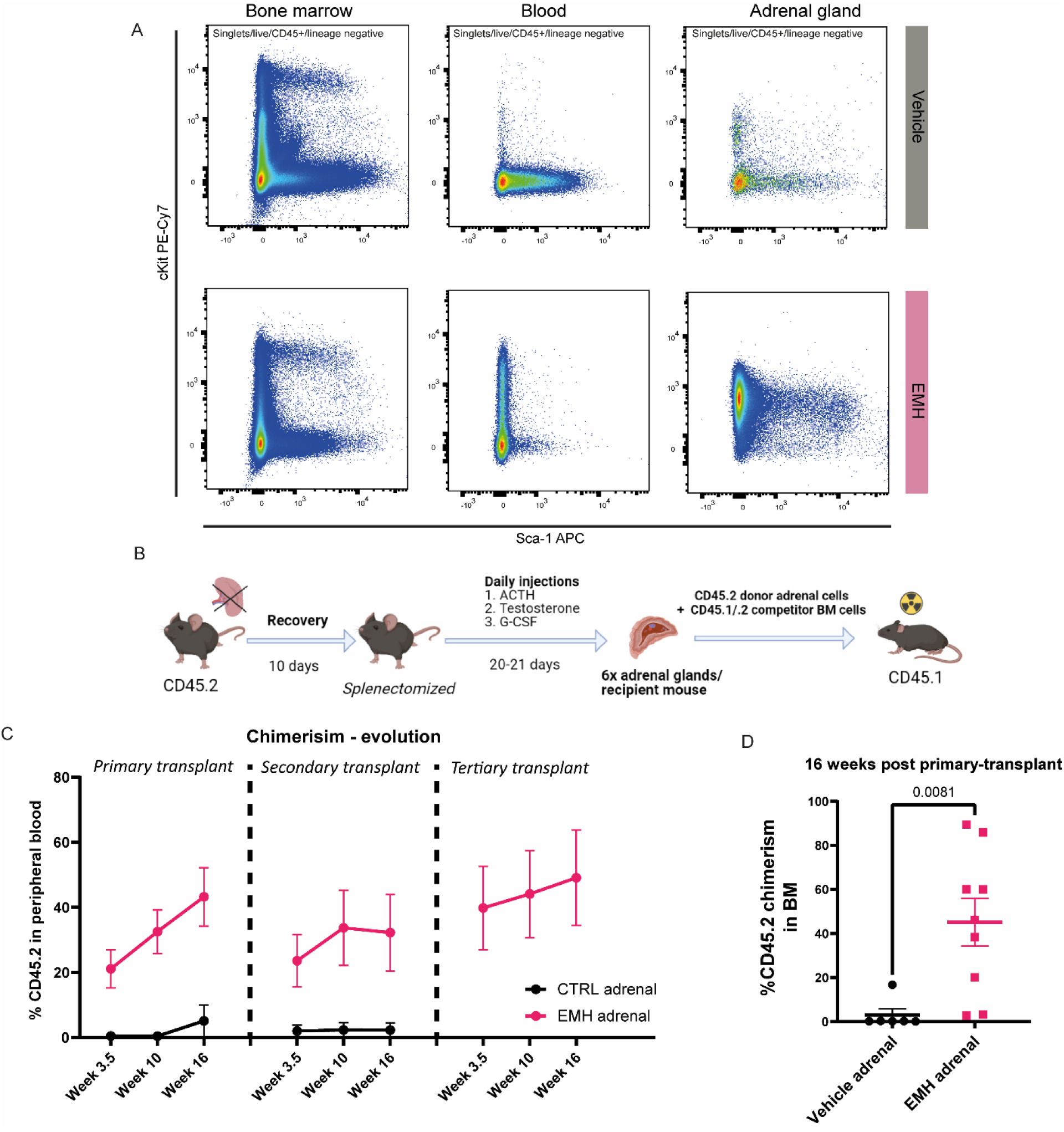
The adrenal gland supports functional, serially transplantable HSPCs. **A** Flow cytometry analysis of the BM, blood and adrenal glands of control versus EMH-treated mice, representative panels gated within the CD45+lineage negative gate are shown (control n=10, EMH n=10 mice, three independent experiments). **B** Experimental design of the competitive transplant. The total content of CD45.2+ cells retrieved from 6 adrenal glands were transplanted into a lethally irradiated CD45.1 recipient together with 125.000 CD45.1.2 total BM cells from a competitor mouse. **C, D** Evolution of the CD45.2 donor engraftment measured in peripheral blood and BM, respectively, by flow cytometry (control n=7, EMH n=8 mice, two independent experiments), Data are shown as mean ± SEM. Differences were assessed using a two-tailed unpaired Student’s *t*-test.

Therefore, to study the functionality of the hematopoietic cells in our EMH-induced adrenal glands, we used competitive BM transplantation assays. Hematopoietic stem cells (HSCs) are defined by the capacity for multi-lineage long-term engraftment in vivo upon serial transplantation (26). CD45, a pan-hematopoietic marker, can be found in two isoforms in C57BL/6 mice, either CD45.1 or CD45.2, which allows for reliable tracking of donor, recipient and competitor-derived populations (27). CD45.2 mice were treated with the EMH induction cocktail, and the cells obtained by enzymatic digestion of the adrenal glands were directly transplanted together with CD45.1/.2 BM competitor cells into lethally irradiated CD45.1 mice. Based on CFU data, 6 adrenal glands contain a similar colony-forming potential as 125.000 total BM cells and, therefore, we transplanted the total cellular content of 6 adrenal glands together with 125.000 total BM competitor cells for a 1:1 competitive transplant assay. CD45.2+ cells in the blood of the CD45.1 primary recipient indicate engraftment originating from our donor mice and thus reveal adrenal resident HSPCs (Figure 2B). We observed a significant CD45.2 engraftment only for mice receiving CD45.2 donor cells from EMH-induced adrenal glands, but not from control adrenal glands. More importantly, CD45.2 cells obtained from EMH-induced adrenal glands were serially transplantable and circulate in the peripheral blood up to at least tertiary transplants (Figure 2C). Long-term CD45.2 engraftment was also observable in the BM (Figure 2D) of mice receiving cells from EMH-induced donors, thereby serving as proof of the presence of functional HSPCs in the EMH-induced adrenal glands.

### Induced adrenal glands recruit circulating HSPCs

After identifying the hematopoietic supporting capacity of the induced adrenal glands, we sought to define the source of the observed HSPCs. Embryologically, the aorta-gonad-mesonephros (AGM) structure gives rise to both the definitive HSPCs and the adrenal cortex. During embryonic development, in the AGM, HSPCs derive from endothelial cells in an endothelial-to-hematopoietic (EHT) transition (28). Even if unlikely, we wanted to exclude that the HSPCs we observed in the adrenal glands could develop *in situ* from non-hematopoietic adrenal cells upon EMH induction. To test this hypothesis, we transplanted splenectomized mice with GFP+ total BM cells to obtain a mouse with a GFP+ hematopoietic system. After recovery post transplantation, mice were treated with the EMH induction cocktail for 20 days (Figure 3A). CFU assays showed the presence of hematopoietic progenitors in the induced adrenal glands (Figure 3B), but not in control mice, as well as in the BM of all mice (Figure 3C). It should be noted that the baseline number of colonies was reduced by about 50% in irradiated mice as compared to all other experiments performed in non-irradiated EMH-induced adrenal glands. Close to 100% of all colonies present in the adrenal glands CFU assays were GFP+, indicating a BM origin of the hematopoietic cells in the adrenal gland upon EMH induction. Overall, these results indicate that HSPCs found in the adrenal gland are recruited from the BM into the induced adrenal gland and do not arise *de novo* in the organ.

**Figure 3.**
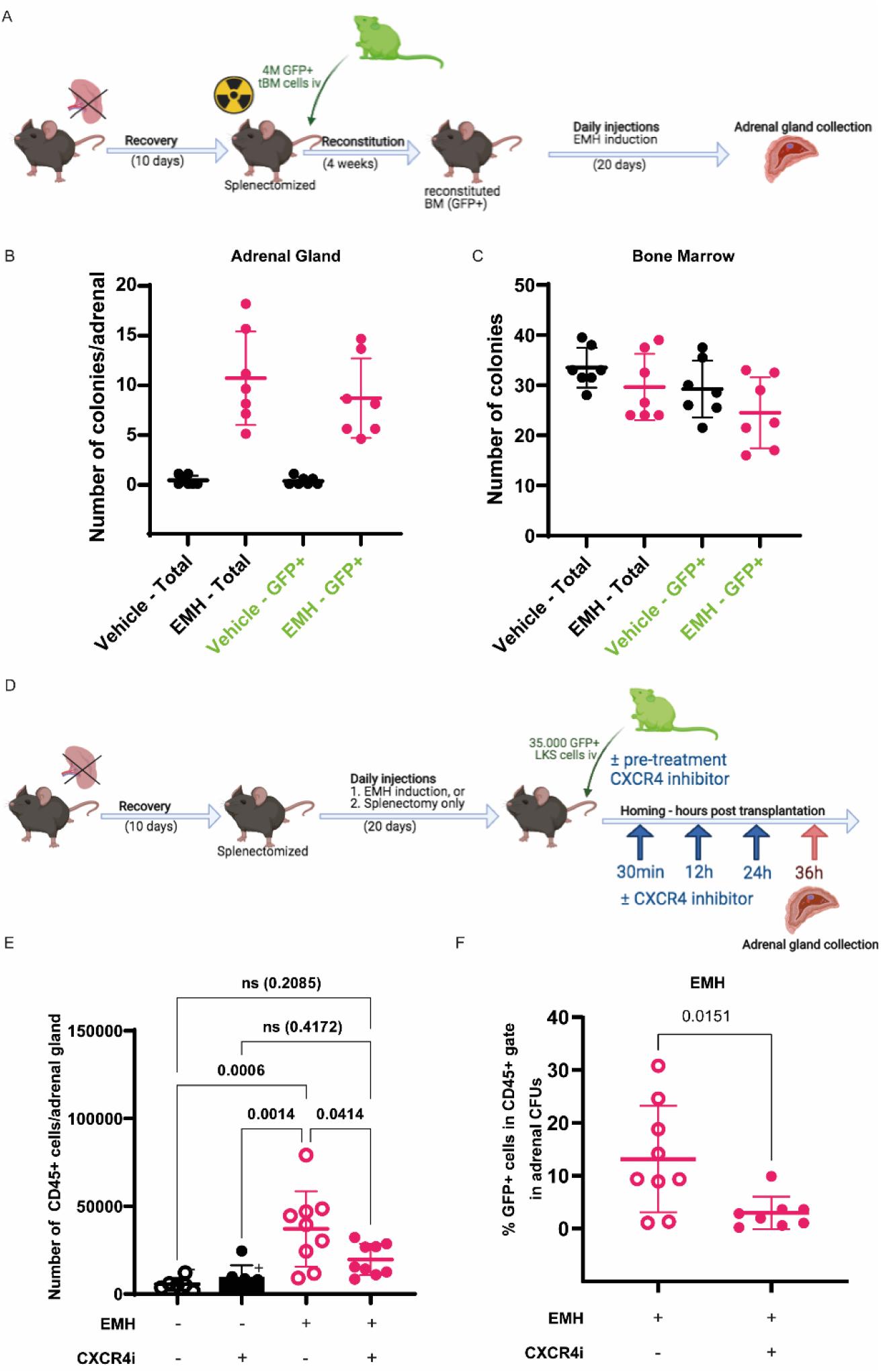
The adrenal stroma recruits and supports circulating hematopoietic progenitors. **A** Experimental design for EMH induction after transplant with GFP+ BM. **B, C** CFU assay, number of total colonies and GFP+ colonies per single adrenal gland (**B**) and 10,000 CD45+ total bone marrow cells (**C**) (n=6 per experimental group, two independent experiments). **D** General experimental design for homing assay. **E** CD45+ cells in EMH-induced adrenals glands retrieved from mice treated with plerixafor (CXCR4i), evaluated by flow cytometry (two independent experiments, n=8 for control groups and n=10 for EMH-induced groups). **F** Percentage of GFP+ cells from the recovered cells grown in methylcellulose CFU culture after completion of the CFU assay (8 days post-plating), quantified by flow cytometry (n=9 control mice and n=8 experimental mice). Data are represented as mean ±SD and groups were compared with one-way ANOVA followed by Tukey’s multiple correction test (**E**) or two-tailed, unpaired Student’s *t*-test (**F**).

### CXCL12 is required for homing and retention of hematopoietic cells in the induced adrenal gland

Once we had determined that the induced adrenal gland can be colonized by hematopoietic cells with CFU potential originating from the BM, we hypothesized that the CXCL12-CXCR4 axis would be involved in this phenomenon, given its crucial role in homing of hematopoietic cell to the BM (29). To evaluate this hypothesis we used plerixafor, a pharmacological antagonist of CXCR4. We performed the EMH-induction protocol and then injected the EMH-induced mice with 35.000 GFP+ LKS cells treated with plerixafor. We administered plerixafor intraperitoneally at 30 min, 12 hours and 24 hours post-LKS injection (29). The mice were sacrificed 36 hours post-LKS injection and the adrenal glands evaluated for CD45+ counts and GFP+ CFU potential (Figure 3D). We expected plerixafor to hamper the colonization of the adrenal niche by the injected GFP+ LKS and therefore decrease the number of GFP+ cells produced in the CFU assay. Congruently, we observed a marked decrease in CD45+ cells in the adrenal glands of induced mice treated with plerixafor (Figure 3E), indicating that the CXCL12-CXCR4 axis is necessary for the retention of CD45+ cells in the adrenal niche. We did not observe differences in the absolute number of colonies formed by plerixafor-treated and non-treated EMH adrenal glands (data now shown). Consistent with our hypothesis, retrieval of the cells from the methylcellulose culture and analysis of their GFP signal by flow cytometry, showed a decrease in the percentage of GFP+ cells that had arisen in the colonies obtained from the organs of the EMH-induced animals treated with plerixafor (Figure 3F), indicating that the CXCL12-CXCR4 axis is required for the homing of hematopoietic cells with CFU potential to the adrenal gland.

Finally, we looked into the presence of CXCL12-abundant reticular (CAR) cells in the adrenal gland, which have been described to be essential for hematopoietic support and retention in the BM (30). For this, we took advantage of a widely characterized CXCL12-GFP knock-in murine reporter model (31). We identified GFP+ cells in both the induced and non-induced adrenal gland by flow cytometry (data not shown) and confirmed our findings with confocal microscopy (Figure 4). These cells were of reticular morphology, reminiscent of that described for CAR cells in the BM (32). Surprisingly, and despite the effect of CXCR4 blockage in induced as compared to non-induced adrenal glands, we observed no obvious differences in CXCL12-GFP+ (CAR) cell numbers or morphology between groups. Taken together, our results show that the adrenal stroma shares immunophenotypic features with the BM stroma and contains CAR-like CXCL12+ cells. Furthermore, our data is compatible with hematopoietic cells being actively retained in the adrenal niche by CXCL12-CXCR4 signaling and not passively sequestered in the organ due to an increase in adrenal vascular volume.

**Figure 4.**
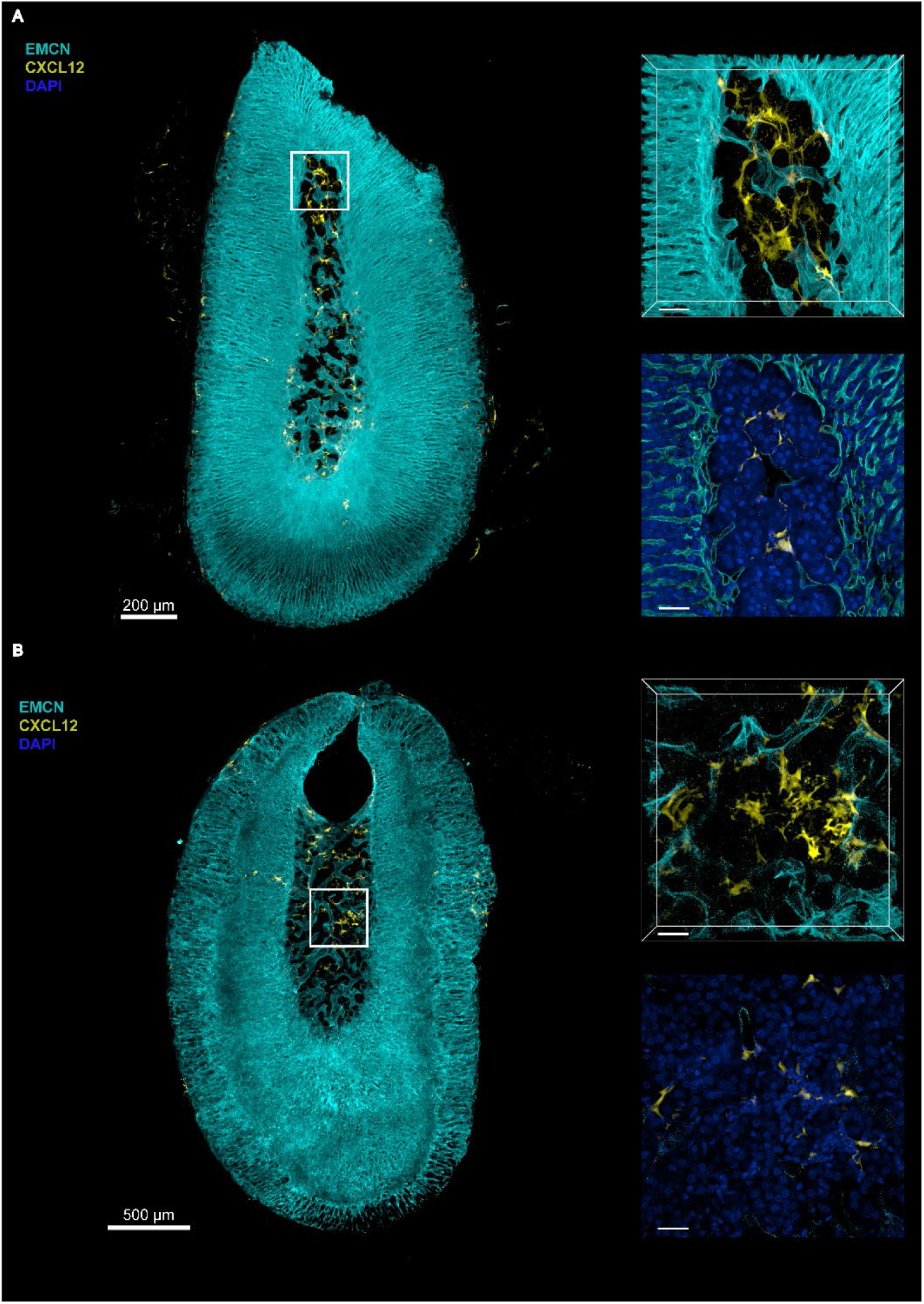
The murine adrenal gland contains CXCL12-positive cells. 3D Microscopy of murine adrenal glands: Representative 3D sections and optical slices of immunostained adrenal glands from (A) control-treated (n=3) and (B) EMH-induced (n=3) Cxcl12^GFP^ transgenic mice showing Endomucin (EMCN;Cyan), CXCL12-GFP (yellow) and 4’,6-diamidino-2-phenylindole (DAPI;blue). Scalebars (A) 200μm, (B) 500μm and cropped sections and optical slices representing 40μm.

### Human myelolipoma is positive for BM stroma markers and contains CXCL12+ cells

Myelolipoma, a benign tumor composed of adipose and hematopoietic tissues, is frequently found in the adrenal gland, particularly in the context of endocrine disorders that associate elevated ACTH levels. We hypothesized that myelolipomas might recapitulate a phenomenon like the one we observe in the adrenal gland of mice treated with our induction cocktail. For this, we retrieved myelolipoma samples originally collected at the Centre Hospitalier Universitaire Vaudois (CHUV), Lausanne, Switzerland. The clinical characteristics of our myelolipoma cohort are summarized in Supplementary Table 1. We then compared our samples with the published registry gathering all reported data cases of adrenal myelolipoma recently published by Decmann et al (33), shown in Table 1. In agreement with published statistics, one patient in our cohort presented an adrenal endocrine disorder (primary hyperaldosteronism). Surprisingly, we also found that 20% of the patients in our cohort had a history of splenectomy whereas the previously published prevalence of splenectomy in the general population is around 0.4% (34). Overall, our myelolipoma cohort is comparable with the literature without noteworthy differences except for the incidence of splenectomy.

**Table 1.** Detailed characteristics of the different immunohistochemical stains on the myelolipoma samples and two controls (healthy BM (J) and adrenal adenoma(G)), quantification represent number of positive cells per samples (-corresponds to no positive cell, + to +++ correspond to linearly increasing number of positive cells)

| Sample ID | CD34 | CD73 | CD90 | CD146 | CD271 | CXCL12 | NESTIN |
| --- | --- | --- | --- | --- | --- | --- | --- |
| **A** | + | / | +++ | +++ | / | ++ | +++ |
| **B** | - | + | - | ++ | + | + | ++ |
| **C** | + | + | + | +++ | +++ | ++ | ++ |
| **D** | - | + | + | ++ | +++ | +++ | + |
| **E** | - | +++ | +++ | +++ | +++ | + | +++ |
| **F** | - | ++ | ++ | +++ | +++ | + | +++ |
| **G** | - | ++ | + | ++ | +++ | - | +++ |
| **H** | - | ++ | + | ++ | +++ | + | +++ |
| **I** | + | +++ | + | ++ | +++ | + | ++ |
| **J** | - | + | + | + | ++ | - | + |
| **K** | + | + | ++ | ++ | +++ | + | +++ |

We performed IHC studies of our myelolipoma samples using a panel of markers designed to evaluate the human BM stroma, namely CD73, CD90, CD146, CD271, CXCL12 and Nestin, as well as CD34 to target hematopoietic progenitors and endothelial cells (35). CD73 and CD90 are two of the positive selection markers described in the minimal criteria for defining multipotent bone marrow stromal cells (36). CD146 has been described to stain perivascular BM stromal cells with multipotent stem cell properties (37, 38). CD271 stains for marrow stromal progenitor cells (37, 39, 40). CXCL12 and Nestin have also been shown to play a role in hematopoietic support within the human BM stromal compartment (41, 42). Using paraffin-embedded samples stored from the diagnostic workup of our patients, we stained consecutive sections and examined the presence of positive cells. We included, as controls, a human BM sample and an adrenal adenoma. CD34 and CD90 were expressed in only a fraction of our samples. CD73, CD146 and CD271 and Nestin were expressed across all samples (Supplementary Figure 2). Notably, CXCL12 was present in all myelolipoma samples (Figure 5 and Table 1) but not in the adrenal adenoma control. Furthermore, CXCL12+ cells were of reticular morphology (Figure 5, inset). Taken together, our results indicate the reproducible detection of stromal cells in myelolipoma samples with similar characteristics as the BM stroma. Thus, our data supports the use of myelolipoma as a surrogate to understand the composition of a minimal supportive hematopoietic niche and mirrors our inducible adrenal niche model.

**Figure 5.**
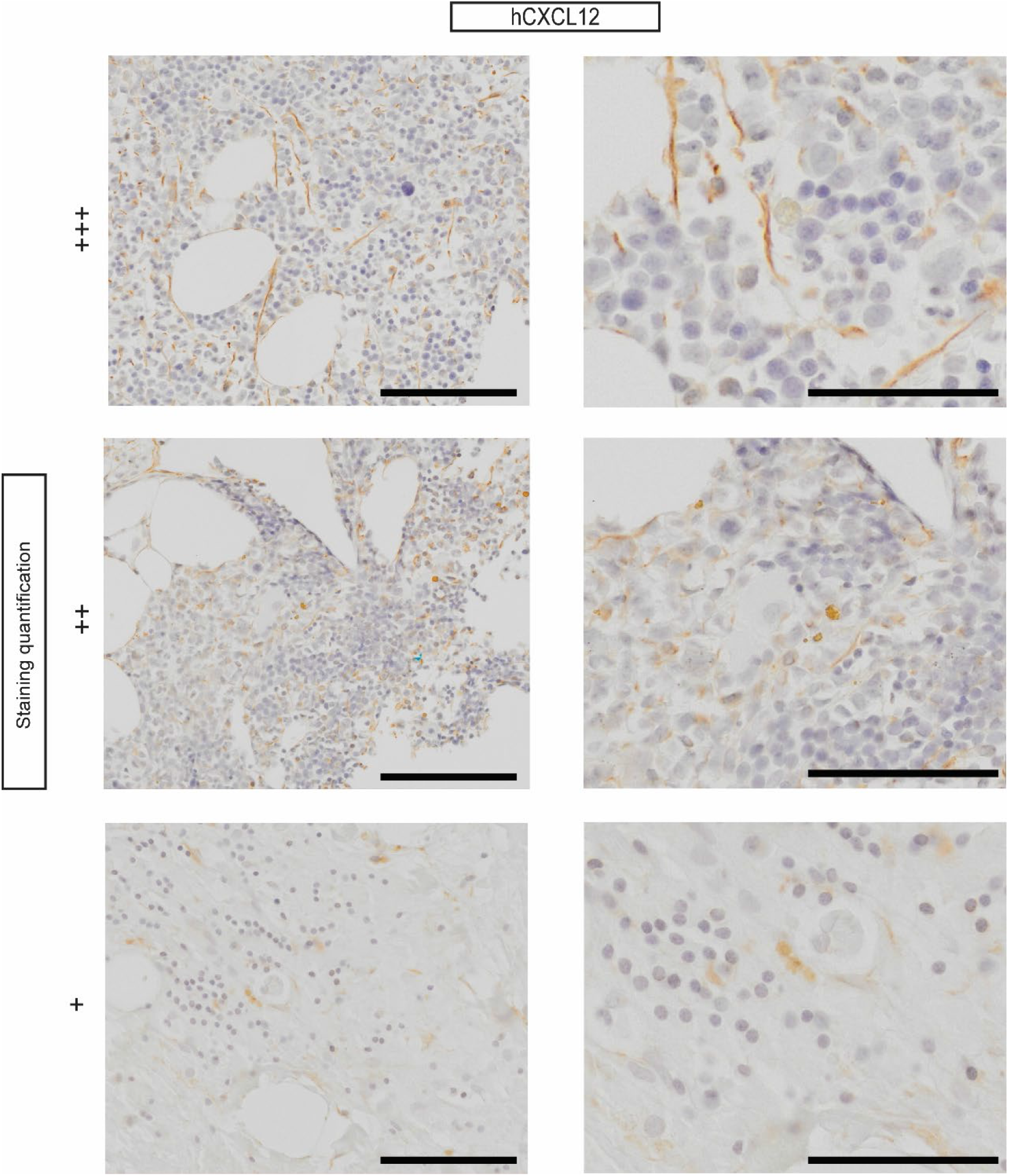
CXCL12+ cells of stromal morphology are present in human myelolipoma. Representative images of CXCL12 IHC stains of human myelolipoma samples corresponding, from top to bottom, to patients D, C and I (see Table 1 for details). Scale bars correspond to 200 μm and 100 μm in the left and right columns, respectively. Dotted stain was considered background and disregarded during assessment.

## Discussion

Here we show that the adrenal gland can be transformed into a hematopoietic supportive environment and used as a model to study the minimal stromal components of a non-ossified *de novo* hematopoietic niche. The EMH-induced adrenal niche contained CXCL12+ cells with classical reticular morphology and supported HSPCs, including serially transplantable stem cells. Our findings are supported by the presence of CAR-like CXCL12+ cells in human adrenal myelolipoma.

The observation that the adult adrenal gland can support hematopoiesis upon hormonal stimulation is particularly interesting as the adrenal cortex originates from the AGM, the structure that gives rise to definitive HSCs during embryonic development. The adrenal gland is therefore ontogenically related to a hematopoietic supportive structure and can be transformed into an adult niche, a phenomenon that has been clearly described in the spleen and the liver (19). Furthermore, it has been recently shown in a human cell atlas of fetal gene expression, that the fetal adrenal gland contains limited erythropoiesis, pointing to the hematopoietic supportive capacity of the adrenal gland (43). Furthermore, the propensity of myelolipoma to develop in the adrenal gland suggests a specific hematopoietic supportive population in this location. Feng et al. (44) put forth a model where myelolipoma originates from a mesenchymal progenitor cell giving rise to the stromal compartment and recruiting hematopoietic cells, something that would be in line with both our observations.

We initially attempted to fully recapitulate myelolipomas in mice with both the adipocytic compartment and hematopoietic cells. Based on historical publications using rats as models, we developed a model using hormonal stimulation in splenectomized mice. G-CSF was used to stimulate the hematopoietic compartment. G-CSF treatment alone directly stimulates HSPCs proliferation and mobilization, accelerating exit of severe neutropenia by an average of 3-6 days in humans (45, 46). In accordance with the Seyle and Stone (24) original description, our cocktail also contained testosterone. The effect of androgens as a stimulant of hematopoiesis has been thoroughly described and is used in patients to treat bone marrow insufficiency, specifically in the context of telomeropathies and Fanconi anemia (47–49). ACTH was used daily to induce EMH in the adrenal gland. The incidence of myelolipoma increases several-fold in patients suffering from congenital adrenal hyperplasia, a disease of the cortisol axis that increases the levels of ACTH (33). Consequently, our induction cocktail is based on stimulation of both the adrenal gland, with ACTH, and the hematopoietic system, with G-CSF and testosterone. We however failed to observe adipocytes in our model, which suggests that mature adipocytes might not be necessary for hematopoietic support in the adrenal EMH.

In our collection of human adrenal myelolipoma samples, patients presented similar characteristics as published in the literature. IHC, using markers described in the BM stroma, consistently showed the presence of cells expressing CXCL12 and Nestin. Both proteins are known to be expressed by BM stromal cells capable of providing hematopoietic support, and have been linked to adrenergic innervation (50). Moreover, the CXCL12/CXCR4 axis is important for solid tumor metastasis (51), which, given the abundance of CXCL12+ cells in the adrenal gland, could explain why this organ is a common site of metastatic spread (52). We noticed a previously unreported increased rate of splenectomy years before the diagnosis of myelolipoma within our cohort of patients (20% in our sample versus 0.4% in the general population). This finding could be explained in several ways. One should be aware that patients who had their spleen removed might be monitored more closely than the general population, and an increase in incidentalome due to more frequent imaging is a possibility (53). Another possible explanation would be that spleen removal induces a change in hematopoietic regulation, therefore inducing a general environment more prone to EMH. The third possibility is a limited sample size which might lead to an artifactual overrepresentation of splenectomies rate in our cohort. Because detailed surgical history was not reported in the previously published systematic review of adrenal myelolipomas (33), a larger cohort of myelolipoma patients would be needed to answer this question. From the perspective of hematopoietic pathophysiology, it would be of great interest to investigate the potential role of splenectomy on EMH. Finally, from a phylogenetic standpoint, examples of boneless, adipose-associated hematopoiesis exist in vertebrate evolution and appear as early as in jawless fish in the form of the dorsal fat body (54)

In conclusion, we present our model as a novel tool to increase our understanding of the physiology of hematopoietic support and to facilitate the development of a boneless niche model. The composition of what constitutes the simplest unit of hematopoietic niche, supporting both HSC self-renewal and progenitor expansion, remains largely unknown. Since the exact composition needed to recapitulate *enough* complexity of the hematopoietic microenvironment for it to be functional is still undefined, a deeper understanding of a minimalistic niche, like the inducible adrenal one we report, has the potential to aid in the development of biomedical and tissue engineering applications.

## Materials and methods

### Animal maintenance and bone marrow transplantation

Mice were housed in 12h day-night light cycles, at 24°C in ventilated cages and provided with sterile food and water. Experiments were carried out in accordance with the Swiss law and with approval of the cantonal authorities (Service Vétérinaire de l’Etat de Vaud). No statistical methods were used to predetermine sample size. Mice were randomly assigned to control and intervention groups. The investigators were not blinded to allocation during experiments and outcome assessment unless otherwise stated.

C57BL/6J (CD45.2) and C57BL/6J Ly5.1 (CD45.1) female mice were purchased from Charles River Laboratories International. Double congenic mice (CD45.1/.2) were bred in-house, by crossing C57BL/6J mice and CD45.1 mice. B6.GFP (C57BL/6-Tg(CAG-EGFP)131Osb/LeySopJ) were bred in-house. CXCL12-GFP knock-in mice (55) were a gift from César Nombela-Arrieta (University of Zurich) and Takashi Nagasawa (Osaka University), and were bred in-house. B6 ACTB-EGFP (Tg(CAG-EGFP)1Osb) mice were bred in-house.

For transplantation assays, recipient mice were irradiated (lethal x-ray irradiation 8.5 Gy, split in two 4.25 Gy doses 4 h apart using an RS-2000 X-ray irradiator (RAD SOURCE). Transplanted cells were administered the day following irradiation via tail-vein injection. For two weeks after lethal irradiation, mice were treated in the drinking water with paracetamol (500mg Dafalgan in 250ml water) and antibiotics in the form of cotrimoxazole (60mg/Kg/day: 250ml of water with 2.5ml of Bactrim (200mg sulfamethoxazole /40mg trimethoprim)).

For assessment of the HSC compartment in the adrenal gland, 8-12-week-old CD45.1 female were irradiated and transplanted the next day with a mix of enzymatically digested adrenal cell suspension containing CD45.2 cells and 125.000 total bone marrow CD45.1/.2 competitor cells.

For secondary and tertiary transplantation experiments, the irradiated recipients were injected intravenously with 4 million total BM cells from the primary recipient or secondary recipients, respectively, suspended in a volume of 300 μl PBS EDTA 1 mM via tail-vein injection. Mice were given at least 18 days of recovery post-transplantation before assessment of the chimerism in peripheral blood.

### Surgery

Thirty minutes before surgery, mice received one subcutaneous injection of 0.1 mg/kg Temgesic/buprenorphine (0.3 mg/mL solution). In addition, mice were given Ibuprofen in drinking water (Algifor Dolo Junior, 2.5 ml of syrup in 250 ml of water, change of water every 2 days) one day prior to surgery and until 3 days post operation. For splenectomy, the animals were anesthetized with isoflurane and placed on a heating pad until re-awakening. Moisture drops (Viscotears) were administered to avoid drying out of the eyes. Isoflurane was used at a concentration of 3-5% for induction of anesthesia and anesthesia was maintained with 2-4% isoflurane. Mice wore a mask to allow constant supply of the anesthetic gas. The animal was placed on its right side and a 1.5 cm incision was made on the left flank skin and then on the peritoneum, caudal to the costal border. The spleen was exposed, and the spleen vessels were ligated with 6.0 resorbable thread (6.0 vicryl, Ethicon) and the spleen was removed using a scalpel blade. The peritoneum first and then the skin were closed with a simple interrupted stitch using 6.0 resorbable thread. After surgery, if needed, blood loss was compensated by intraperitoneal injection of 0.5 mL warm saline solution (0.9% NaCl). A minimal period of seven days was observed before starting any induction experiment.

### EMH induction cocktail and inhibitors

For induction of EMH in the adrenal gland, mice were injected daily for 20-21 days with G-CSF (150 μg/kg filgrastim Neupogen 30mio U/0.5ml Amgen, diluted in Glucosum Bichsel solution 5 % for a final volume of 200ul intraperitoneally on alternate sides of the abdomen), testosterone (310 μg/mouse, NEBIDO 1000 mg/4 ml Bayer diluted in corn oil sigma C8267-500ml for a final volume of 30μl subcutaneously) and ACTH (tetracosactide 20 μg/mouse subcutaneously combined with testosterone or corn oil vehicle - Synacthen Depot susp injectable 1 mg/ml Alfasigma). Stock solutions were kept at 4°C, working stocks were prepared once a week and kept at 4°C.

### Complete blood counts

Blood was collected from the tail vein into EDTA-coated tubes (Microvette® 100 K3 EDTA, Sarsted). The samples were analyzed, and the blood cell counts recorded using the Scil Vet ABC hematology analyzer (Scil, USA).

### Adrenal cell isolation

Adrenal glands were isolated and placed in ice-cold PBS. For preparation of a single cell suspension, adrenal glands were placed in a 15 ml tube containing 4.75 ml of Hanks’ Balanced Salt Solution and were cut open with a sterile scalpel. 250 μl of collagenase I 4% (Gibco, cat: 17018029) was added and falcon tubes were sealed with parafilm and incubated for 30 min at 37 °C with agitation. After incubation, digestion was quenched by dilution with PBS EDTA 1 mM at 4 °C. Cells and remaining adrenal pieces were filtered and smashed on a 100-μm mesh basket filter with a syringe plunger and the filter washed with ice cold PBS EDTA 1 mM. The cells were then centrifuged at 300 g for 7 min at 4 °C and resuspended in 300 μl of PBS EDTA 1 mM and kept on ice until further processing.

### Flow cytometry

Data were recorded using BD LSR-Fortessa (Becton Dickinson) or Gallios (Beckman Coulter) flow cytometers and analyzed using FlowJo (version 10.7.1). The following antibodies were used to assess post-transplant BM reconstitution: CD45.1 FITC (A20, BioLegend 110706; 1:200), CD45.2 Pacific blue (104, BioLegend 109820; 1:200), CD3 PE (17A2, BioLegend 100206; 1:200), CD19 PE (6D5, BioLegend 115507, 1:500), Gr1 APC (RB6-8C5, BioLegend 108412; 1:1000), F4/80 APC (BM8, BioLegend 123116; 1:750).

For BM analysis, we used a biotinylated lineage antibody cocktail against CD3, CD11b, CD45R/B220, Ly-6G, Ly-6C and TER-119 (51-9000794, BD Biosciences; 1:30), Streptavidin-TxRed (S872, Invitrogen; 1:200), Streptavidin Pacific orange (S32365, Invitrogen; 1:100), cKit PE-Cy7 (2B8, BioLegend 105814; 1:200), Sca1 BV711 (D7, BioLegend 108131; 1:150), CD45.2 AF700 (104, BioLegend 109822; 1:100), CD45.1 FITC (A20, BioLegend 110706; 1:200).

### Colony forming unit assay (CFU)

For CFU assays from BM cells, the pelvis, femora and tibias were isolated and cleaned from muscle, tendons and connective tissue using a gauze sponge. Bones were crushed using a mortar and pestle in PBS EDTA 1 mM and filtered with a 70-μm mesh and centrifugated 10 min at 300 g at 4 °C. The pellets were resuspended in 1 mL ice-cold PBS EDTA 1 mM and 10 μl of the resuspension added to an antibody mix containing antibodies against CD45-APC-eFluor 780 at a final concentration of 1:200, Ter119-Alexa Fluor 700 at 1:200 and 10 μl CountBright beads (C36950, Invitrogen). After staining for 30 minutes on ice, the volume was increased to a final volume of 250 μl with PBS EDTA 1 mM containing DAPI at 1 ng/mL and 2 % FBS. The samples were then run on a LSRII (Becton Dickinson, USA) flow cytometer and hematopoietic cells were counted based on FSC/SSC singlets and DAPI-/Ter119-CD45+ as compared to beads to determine the volume of unstained cells for subsequent plating into methylcellulose.

To assess hematopoietic progenitor function via CFU assay, the equivalent of 10.000 DAPI-/CD45+ BM cells in 250 ul of PBS or number of cells equivalent to one third of one adrenal gland, as defined by one third of the final volume of the adrenal single cell suspension, were added to 2.5 ml methycellulose (Methocult GF M3434, STEMCELL Technologies) and 1.1 mL was plated per well in a SmartDish 6-well plate (27371, STEMCELL Technologies, USA) in duplicates. Finally, and after 8 days of culture, the wells were imaged using the STEMvision automated CFU counter (STEMCELL Technologies, USA). Total colony numbers were automatically counted, and manual verification of colony selection was performed and adjusted if necessary. Plots are normalized per adrenal gland.

### Homing assay CXCR4i

After the EMH-induction protocol described above, we FACS-sorted 35.000 GFP+ LKS cells per recipient mouse into PBS plus 2% FBS. We incubated the GFP+ LKS cells with plerixafor 1 μM (AMD3100, Selleckchem) for 15 minutes on ice (56) and injected them intravenously into the recipient mice. Control mice were transplanted with GFP+ LKS cells incubated in PBS only. At 30 min, 12 and 24 hours post-LKS injection, as previously described (29), we administered plerixafor dissolved in PBS at a dose of 10 mg/kg intraperitoneally to recipient mice. Control mice were injected analogously with the vehicle. At 36 hours post-LKS injection, the mice were sacrificed, and the adrenal glands evaluated for CD45+ numbers and GFP+ CFU potential as described above.

### Histology

For the histological assessment of BM, the extracted murine femora and tibias were fixed in 10 % formalin for 24h at RT and washed 3 times for 15 min in PBS at RT. The bones were then decalcified in EDTA (Sigma Aldrich 607-429-00-8) 0.4 M pH 7.2 for 3 weeks, and then thoroughly washed under running tap water. The bones were re-fixed in 10% formalin for 5 hours and washed again in tap water for 30 minutes. The bones were placed in 70% ethanol at 4 °C before stepwise dehydration embedding in paraffin blocks. The bones were then cut longitudinally in 3-4 μm sections and stained with H&E on a Tissue-Tek Prisma automate (Sakura) or immunohistochemical staining were performed.

For IHC, detection of rabbit anti-GFP (Abcam, ab6673,), rabbit anti-vWF (Abcam, ab9378,) or rabbit anti-CD45 (eBioscience, 30F-11) was performed manually. After quenching with 3% H_2_O_2_ in PBS 1x for 10 min, a heat pretreatment using 0.1 M Tri-Na citrate pH 6 was applied at 60 °C in a water bath overnight. Primary antibodies were incubated overnight at 4 °C. After incubation of Immpress HRP (Ready to use, Vector Laboratories), revelation was performed with DAB (3,3’-Diaminobenzidine, D5905, Sigma-Aldrich). Sections were counterstained with Harris hematoxylin and permanently mounted. All slides were imaged on an automated slide scanner (Olympus VS120-SL) at 20x or 40x.

### Immunofluorescence and confocal microscopy

Harvested adrenal glands were fixed with 4% paraformaldehyde overnight and subsequently dehydrated with 30% sucrose in PBS for 48-72 hours. Glands were snap frozen in optimal cutting temperature (OCT) compound for long-term preservation. Samples were then thawed and embedded in low-melting temperature agarose and sectioned using a vibratome to generate 200 μm-thick slices, which were blocked overnight in 0.2% Triton X100/1% BSA/10% donkey serum/PBS. Slices were immunostained with anti-Endomucin [Invitrogen Rat anti-Endomucin (V.7C7), and anti-Rabbit Full-Length GFP Polyclonal Antibody (Living colours)] for 2 days in blocking solution, washed overnight with 0.2% Triton X-100, and stained with secondary DyLight 680 donkey anti-rat immunoglobulin G(IgG), DyLight 488 donkey anti-Rabbit tIgG (Invitrogen), and 4’,6-diamidino-2-phenylindole (DAPI, Life Technologies). Whole-mount stained slices were then washed in PBS and incubated overnight in RapiClear 1.52. For observation under the confocal microscope, slices were directly mounted on glass slides while embedded in RapiClear. Confocal microscopy was performed with a Leica STELLARIS V, equipped with a 405-nm laser and a white light laser, using a 20× objective. Confocal image stacks were processed and rendered into three-dimensional volume with Imaris Software (Oxford Instruments, UK).

### Myelolipoma samples

Clinical data was manually extracted from the patients’ medical record. Biological samples, in the form of unstained slides of 5 μm paraffin-embedded sections, were provided by the Institut de Pathologie, CHUV, Lausanne, Switzerland.

Automated immunohistochemistry was performed on Ventana Discovery ULTRA with the antibodies listed below with a Harris counterstain. Slides were imaged on an automated slide scanner (Olympus VS120-SL) at 20x. Quantification was performed by a trained pathologist (S.G.) and an MD-PhD student (F.S.) in parallel based on number of positive cells per sample.

| **Antibody** | **Titration** | **Reference** |
| --- | --- | --- |
| CD90 | 1/100 | Abcam [EPR3132] ab92574 |
| CD105 | 1/900 | Abcam [EPR10145-12] ab169545 |
| CD146 | 1/250 | Abcam [EPR3208] ab75769 |
| CD271 | 1/100 | Abcam [EP1039Y] ab52987 |
| CXCL12 | 1/15 | Thermofisher 79018 |
| Nestin | 1/3000 | Abcam [EPR13012] ab 156571 |
| CD34 | 1//50 | Agilent [QBEnd 10] GA632 |
| CD73 | 1//50 | Cell signaling technology [D7F9A] 13160 |

### Quantification and statistics

p-values were calculated using unpaired two-tailed Student’s t-test or one-way ANOVA with GraphPad Prism 9. To highlight statistical significances in figures, the following annotations for p values were used: *, p-value<0.05; **, p-value<0.01; ***, p-value<0.001; ****, p-value<0.0001.

### Study approval

All procedures were in accordance with the ethical standards of the responsible committee on human experimentation and in accordance with the 1975 Helsinki declaration as revised in 2008. The local ethical commission approved the study (CER-VD, Lausanne, Switzerland) and a specific consent for this study was obtained in cases where a general consent for research was not already available. For cases where the effort to obtain a specific consent were disproportionate, specific consent was waived by the CER-VD according to the provision of the Swiss Federal Human Research Ordinance (HRO, RS 810.301).

## Authors contribution

ON and FS conceived the ideas and obtained funding for the project. FS performed all the experiments with the help of AAC, SFL and AO. FS, ON and AAC analyzed the results and wrote the manuscript. AV and CNA performed and analyzed all confocal microscopy imaging presented in this manuscript. RS and LdL set up the immunostaining panel for myelolipoma. SG assessed the myelolipoma images. First co-authorship was granted based on the final contribution to the completion of the project; FS developed the experimental pipelines and executed the experiments while AAC assisted during the experiments and was involved in the preparation of the manuscript, analysis of the data and finalization of the project.

## Acknowledgement

FS was funded by the SNSF MD-PhD grant 183986. ON was funded by the SNSF Professorship grant PP00P3_144857 and 183725 and, together with AAC, by the University of Lausanne (UNIL). We wish to thank Josefine Tratwal, Paolo Bianco, Mukul Girotra, Shanti Rojas-Sutterlin and Markus Manz for help in the inception of the project, and for providing critical advice along its development. This project would not have been possible without the support of the EPFL animal facility (CPG) and the animal caretakers, in particular Laetita Cagna, Margaux Mouchet and Pierre Dodane. We thank the EPFL and UNIL flow cytometry facilities, in particular Danny Labes, the EPFL histology facility, in particular Jessica Sordet-Dessimoz, and the EPFL Biomolecular Screening Facility. We thank Dr. Nathalie Piazzon (Institut de Pathologie Biobank, UNIL/CHUV, Lausanne) for facilitating the access to human myelolipoma samples. We thank Dr. Takashi Nagasawa, Graduate School of Frontier Biosciences and Graduate School of Medicine, Osaka University, Japan, for the generous gift of the CXCL12/GFP knock-in mice

