## Supplemental information for "Adrenal extramedullary hematopoiesis as an inducible model of the adult hematopoietic niche"

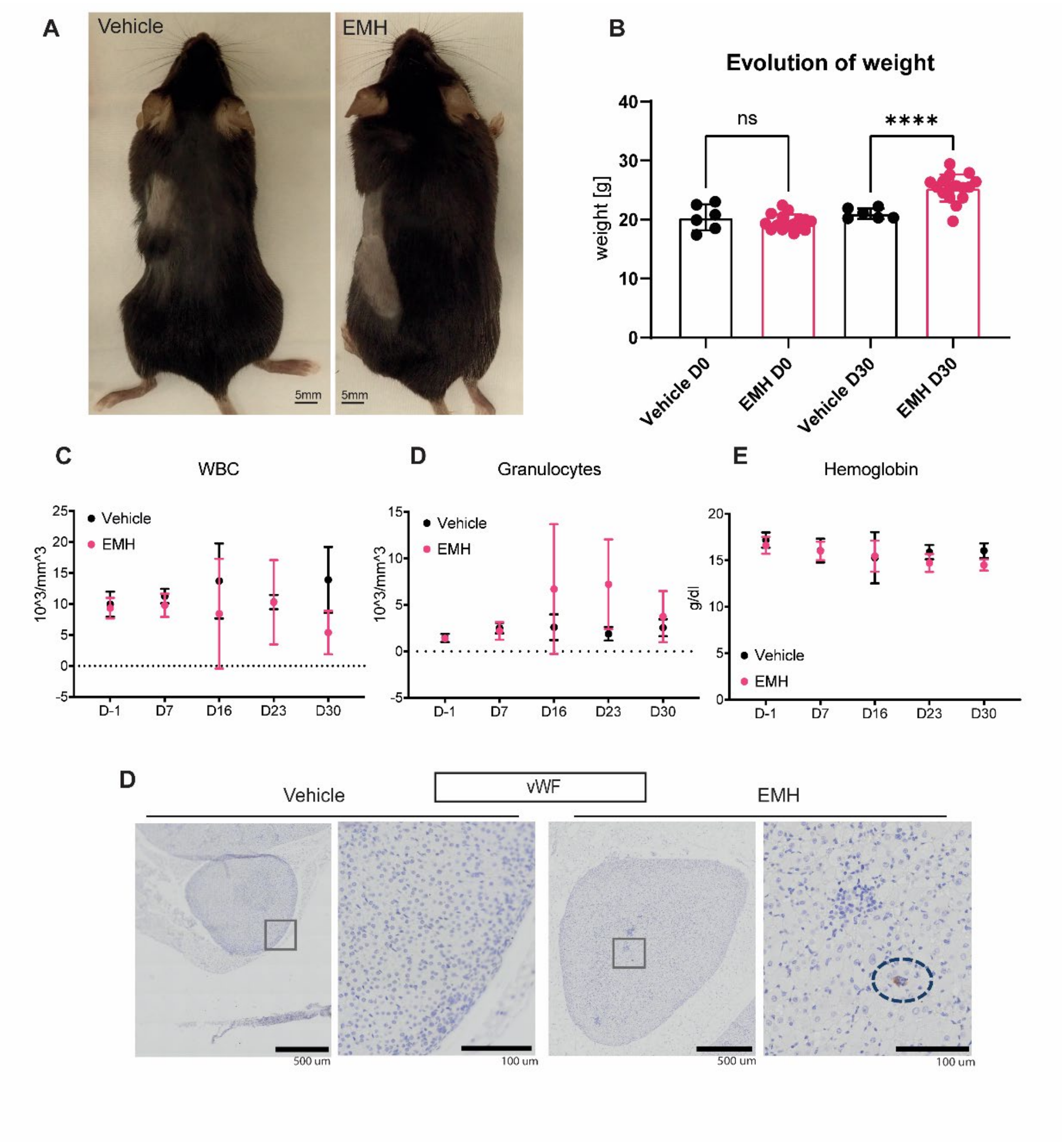


**Supplementary Figure 1**.

A. Representative images of control and EMH mouse after splenectomy and 20 days of daily injections, B. Weight before and after EMH induction, C-E. Complete blood counts during EMH induction, Day 0 represents the day of splenectomy, and D7 the timepoint post-splenectomy before the start of the EMH hormonal induction (n=12 mice per group, three independent experiments). Data are represented as mean ±SD. Unpaired t-test, * p-value<0.05; ** p-value<0.01; *** p-value<0.001; p-value <0.0001.

| Characteristic | Adrenal myelolipoma: a comprehensive review (33) | CHUV cases (1999-2018) |
|---|---|---|
| **Cases** | 440 | 10 |
| **Patient characteristics** | | |
| Age - yr | 51 | 65.1 |
| Gender | No difference | No difference |
| **Tumor characteristics** | | |
| Localization | Adrenal gland only | 6 (60%) adrenal, 4 (40%) pelvic |
| Size | 10.2 cm (min 4 cm, max 430 cm) | 9.1 cm (min 0.3 cm, max 17 cm) |
| Side | 60% right sided | No difference |
| Imaging - calcification | Often | None |
| **Medical history** | | |
| ***Endocrine*** | | |
| Diabetes (type 2) | 41 (9.3%) | 2 (20%) |
| Congenital adrenal hyperplasia | 44 (10%) | 0 |
| Hypercortisolism | 9 (2%) | 0 |
| Primary aldosteronism | 5 (1.1%) | 1 (10%) |
| Androgen secretion | 2 (0.45%) | 0 |
| Hypogonadism | 2 (0.45%) | 0 |
| Primary hyperparathyroidism | 2 (0.45%) | 0 |
| Pheochromocytoma | 2 (0.45%) | 0 |
| ***Hematological diseases*** | | |
| Thalassemia | 12 (2.72%) | 0 |
| Sickle cell diasease | 2 (0.45%) | 0 |
| ***Oncological diseases*** | | |
| Renal cancer | 6 (1.4%), | 0 |
| Prostate cancer | 3 (0.7%) | 0 |
| Breast cancer | 2 (0.45%) | 1 (10%) |
| Gastric cancer | 2 (0.45%) | 0 |
| Endometrial Cancer | 2 (0.45%) | 0 |
| Lung cancer | 0 | 1 (10%) |
| ***Other*** | | |
| Hypertension | 98 (22.3%) | 7 (70%) |
| Atrial fibrilation | 2 (0.45%) | 0 |
| Polytrauma | 0 | 3 (30%) |
| Splenectomy | 0 | 2 (20%) |
| **Laboratory values** | | |
| Blood counts | No information | |
| Anemia | No information | 3 (30%) |
| Lymphopenia | No information | 1 (10%) |
| Lymphocytosis | No information | 1 (10%) |
| Thrombocytosis | No information | 1 (10%) |
| Thrombocytopenia | No information | 1 (10%) |
| Monocytosis | No information | 2 (20%) |
| **Endocrinology workup** | | |
| Hyperaldosteronism | No information | 1 (10%) |

**Supplementary table 1**.

Comparison of the characteristics of patients with myelolipoma from the literature (33) and the collection of samples retrieved from our local hospital.

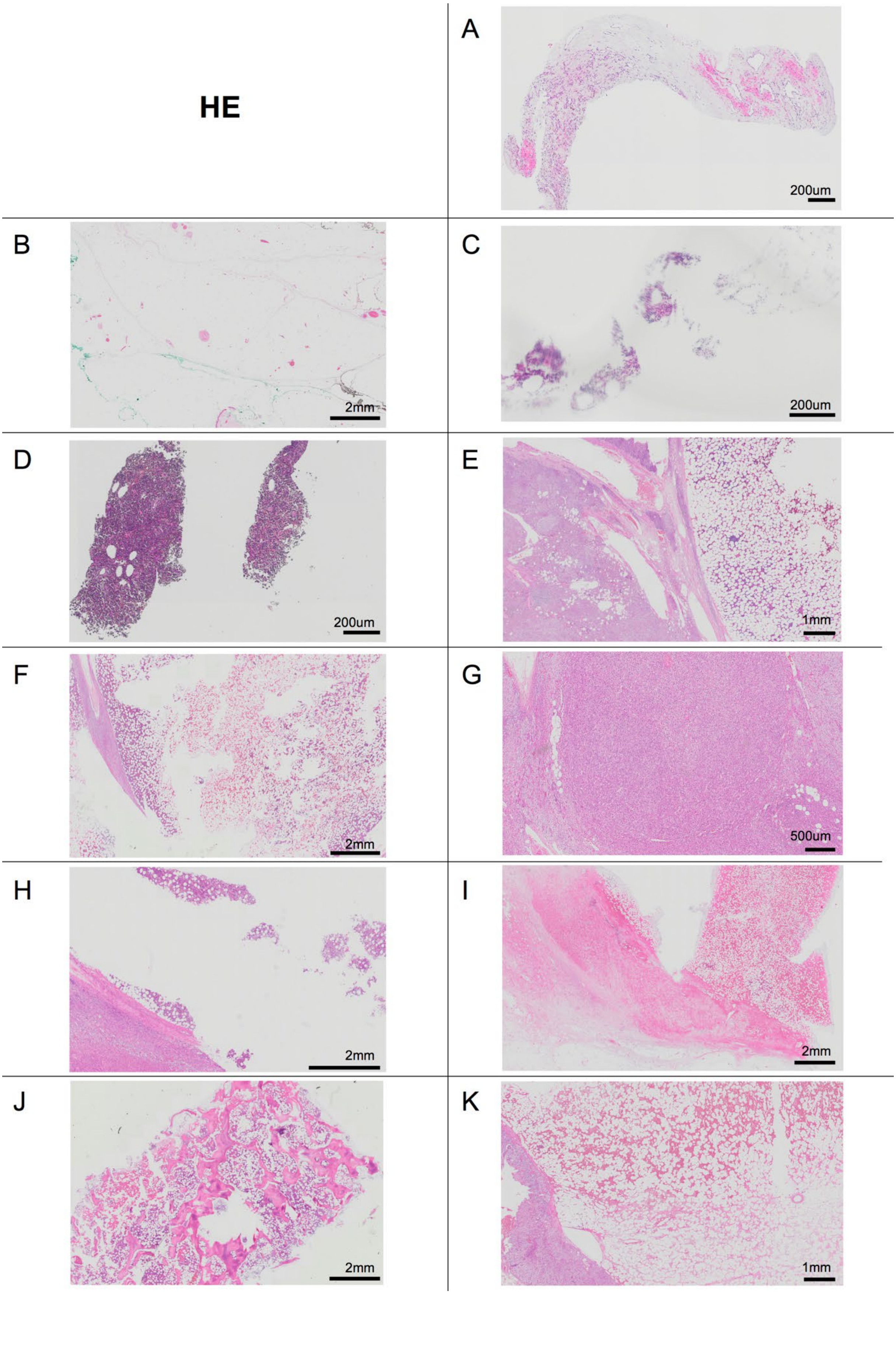
HE
A
200um
B
2mm
C
200um
D
200um
E
1mm
F
2mm
G
500um
H
2mm
I
2mm
J
2mm
K
1mm

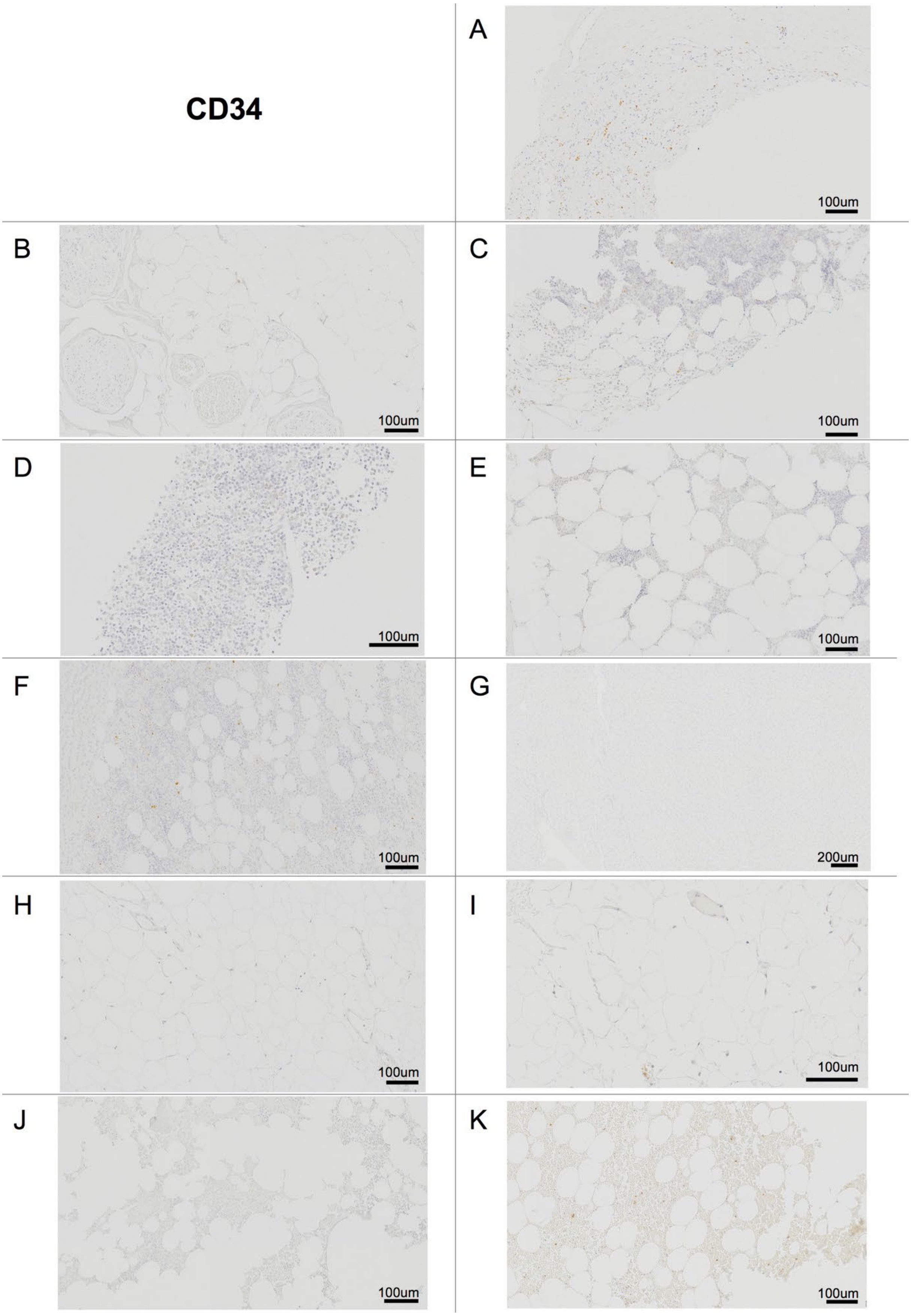
CD34
A
100um
B
100um
C
100um
D
100um
E
100um
F
100um
G
200um
H
100um
I
100um
J
100um
K
100um

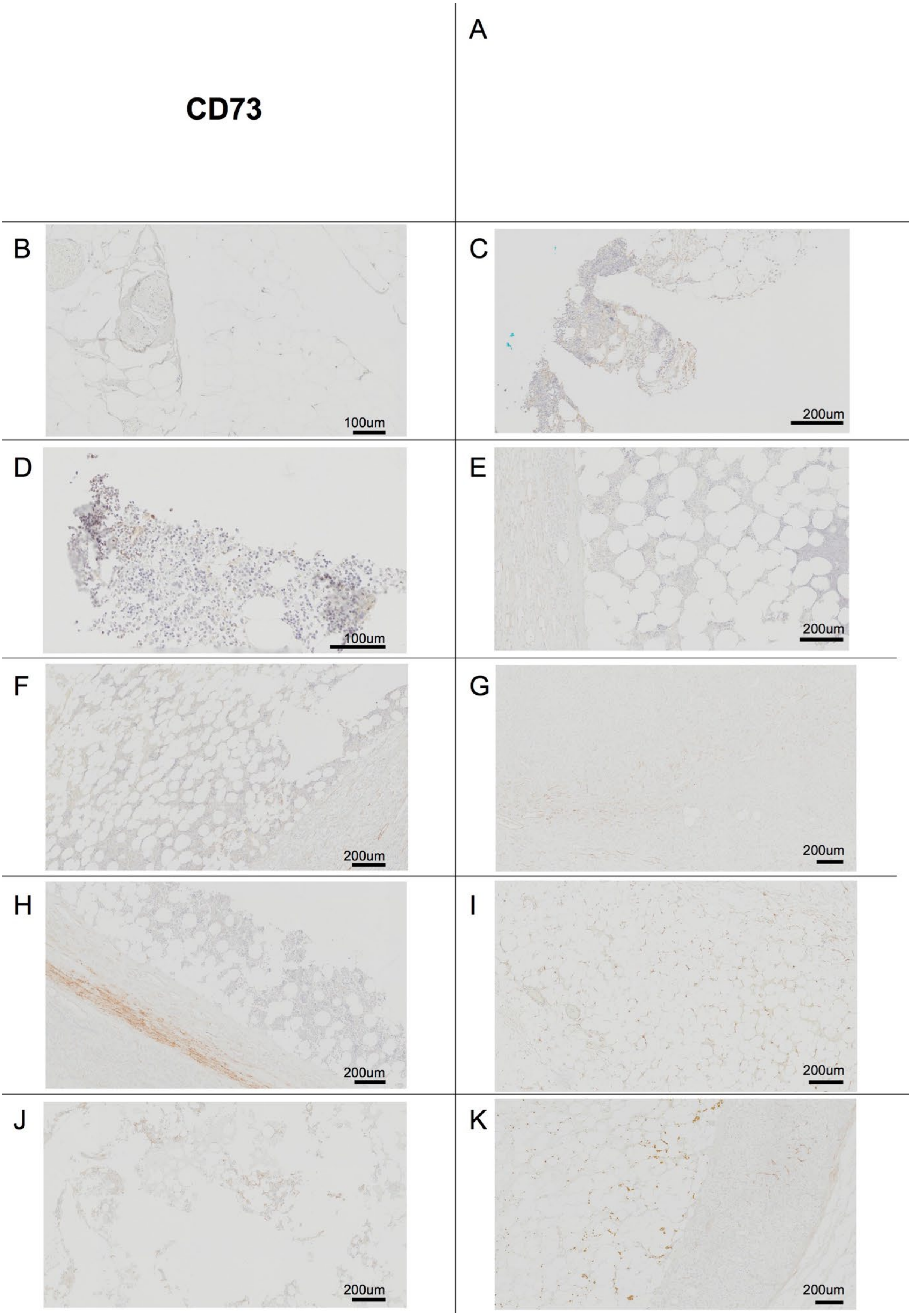
CD73
A
B
100um
C
200um
D
100um
E
200um
F
200um
G
200um
H
200um
I
200um
J
200um
K
200um

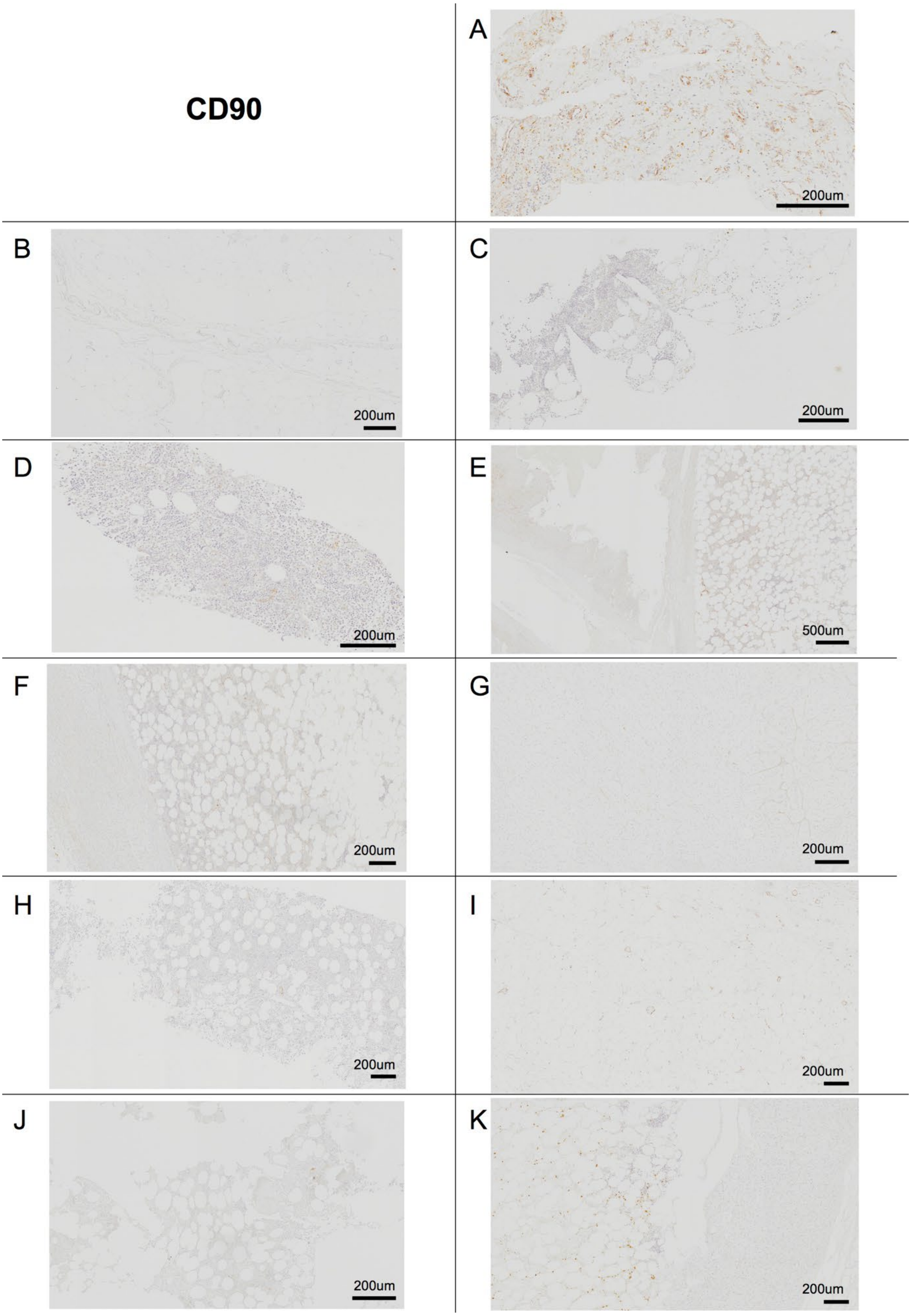
CD90
A
200um
B
200um
C
200um
D
200um
E
500um
F
200um
G
200um
H
200um
I
200um
J
200um
K
200um

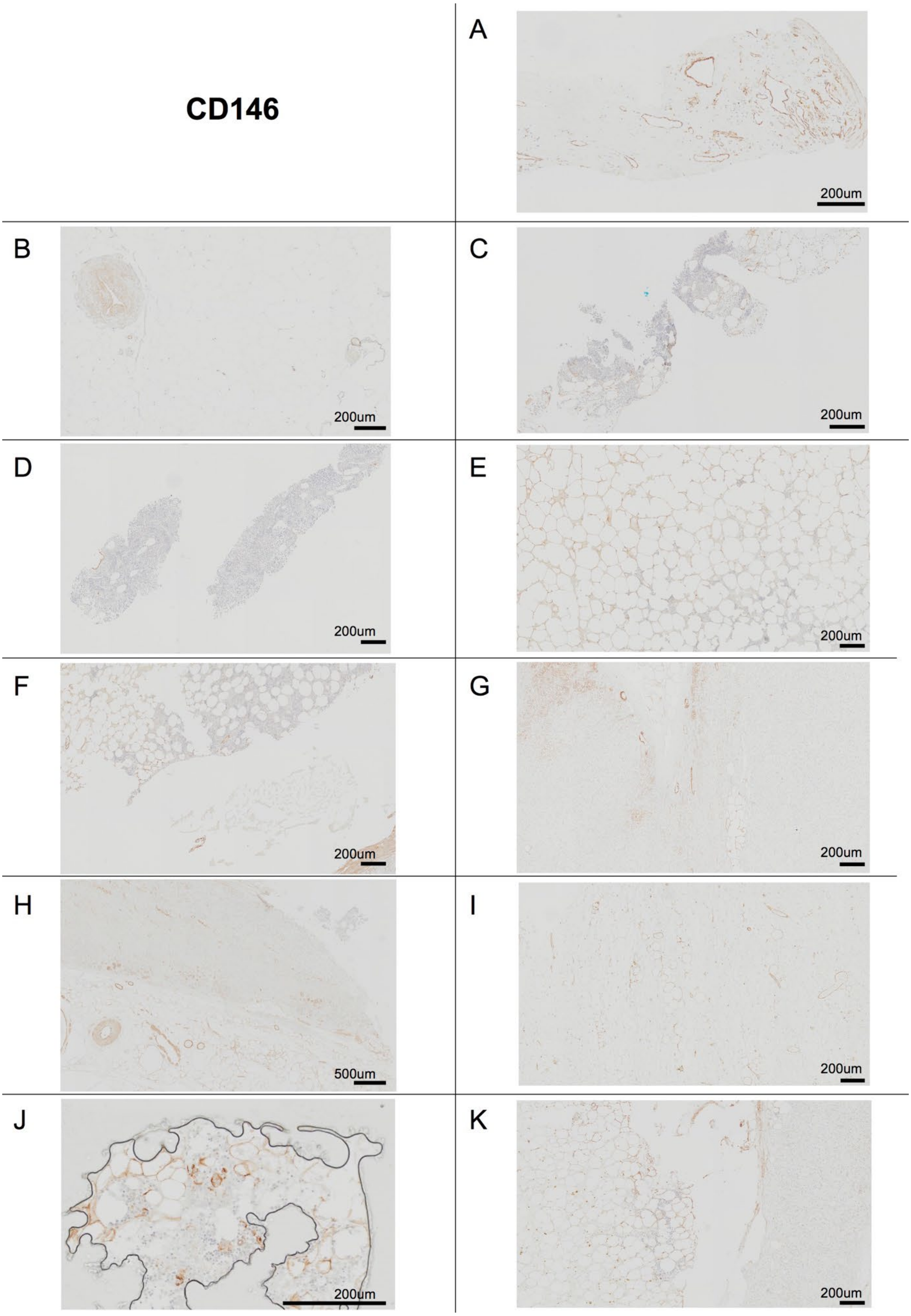
CD146
A
200um
B
200um
C
200um
D
200um
E
200um
F
200um
G
200um
H
500um
I
200um
J
200um
K
200um

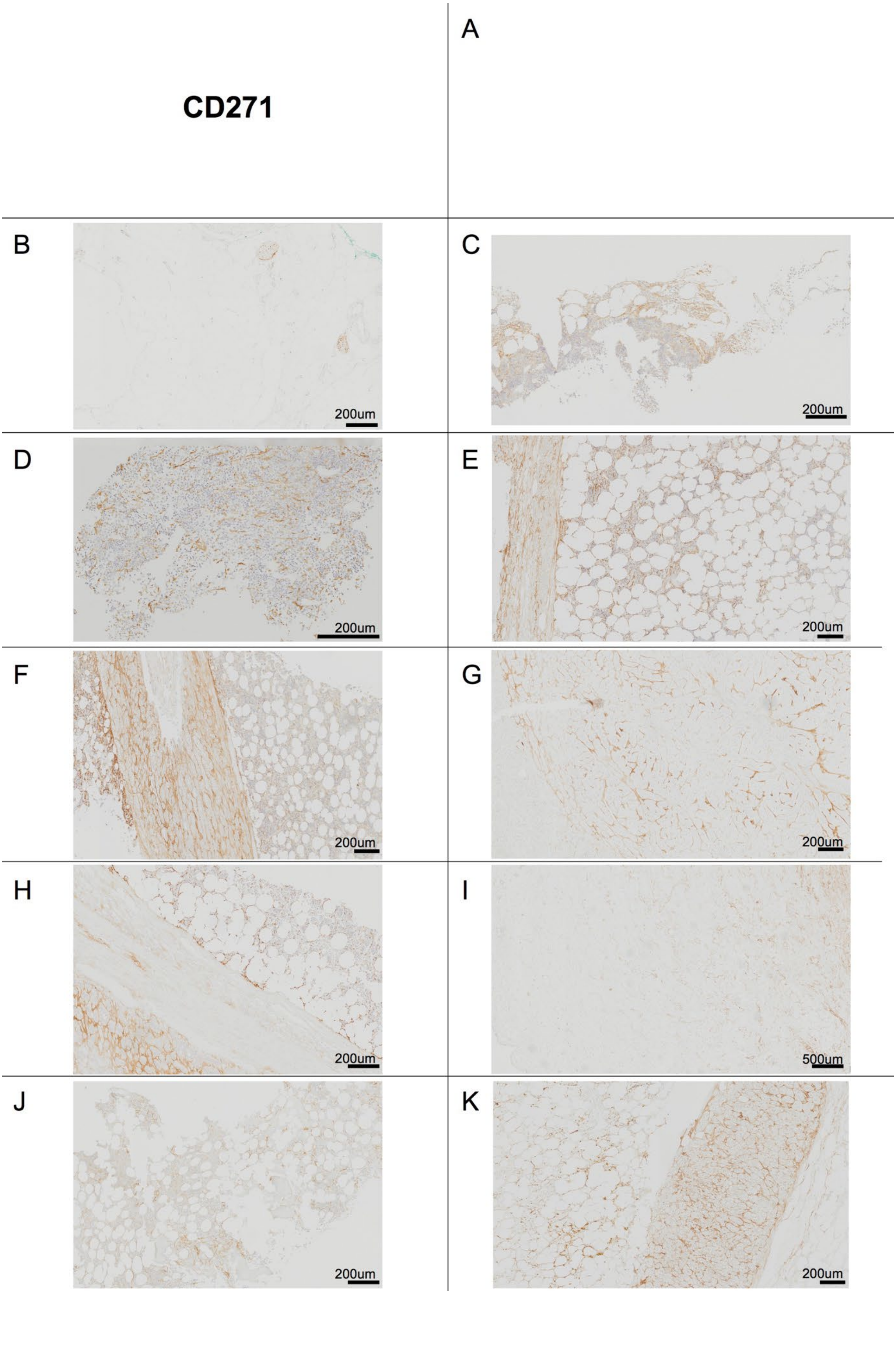
CD271
A
B
200um
C
200um
D
200um
E
200um
F
200um
G
200um
H
200um
I
500um
J
200um
K
200um

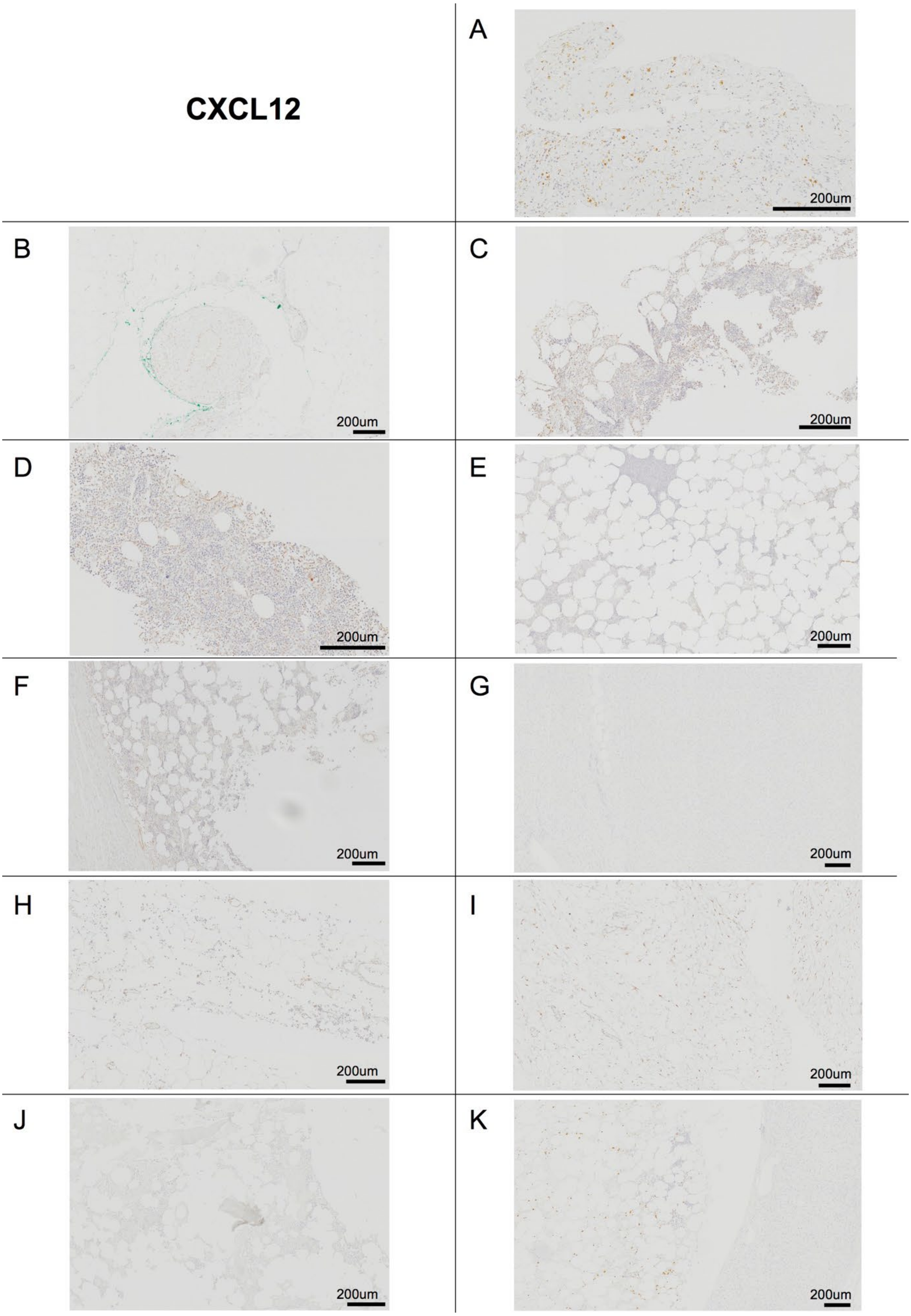
CXCL12
A
200um
B
200um
C
200um
D
200um
E
200um
F
200um
G
200um
H
200um
I
200um
J
200um
K
200um

**Nestin**

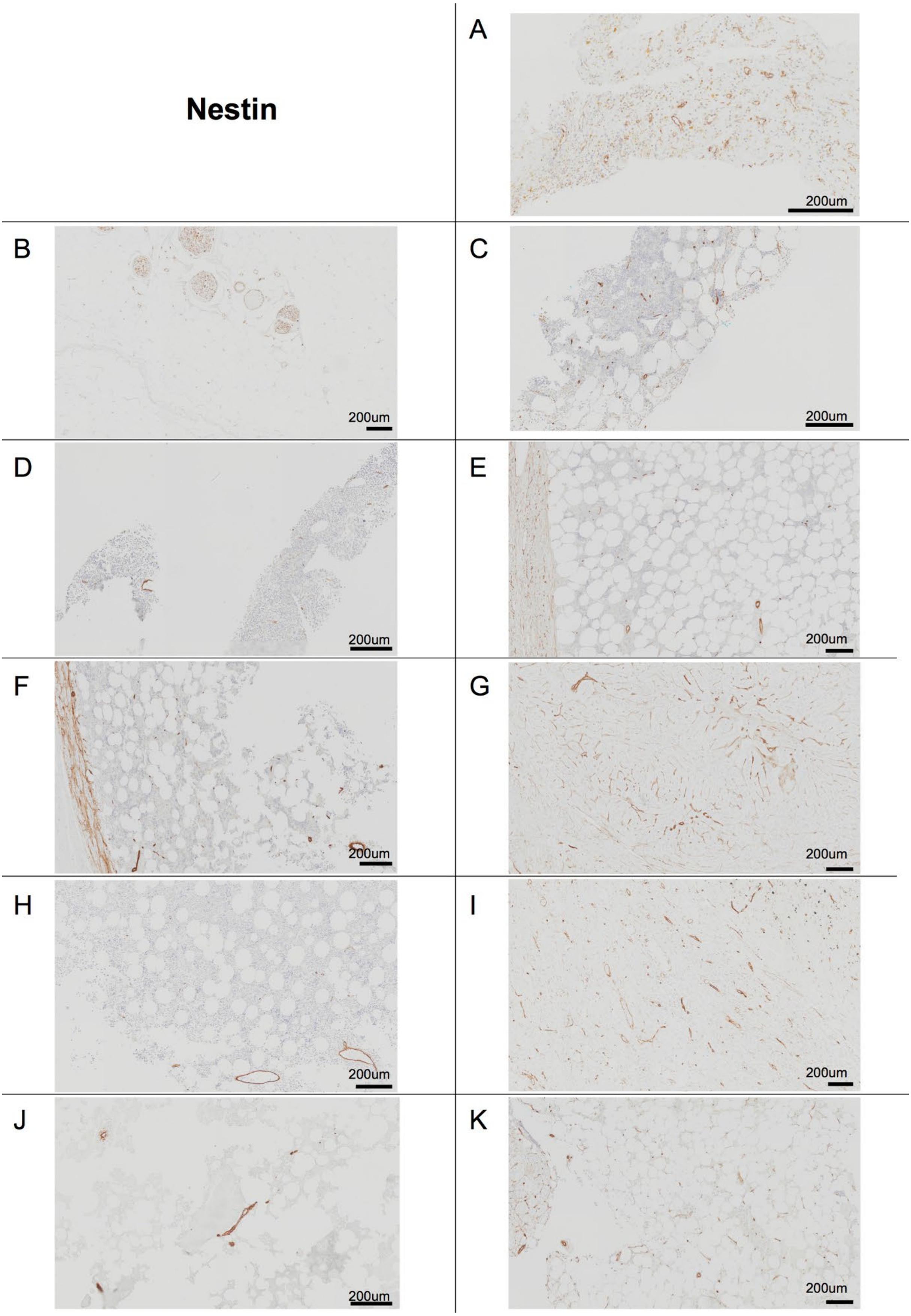

A
200um
B
200um
C
200um
D
200um
E
200um
F
200um
G
200um
H
200um
I
200um
J
200um
K
200um

**Supplementary Figure 2**

Representative images of the different immunohistochemistry stains for each patient myelolipoma. (G) correspond to an adrenal adenoma and (J) to a healthy bone marrow sample, remaining panels are myelolipoma samples. Specific stains or antibodies are indicated on the top left.
